# Dysregulation of U12-Type Splicing in Lupus Neutrophils

**DOI:** 10.1101/2025.11.06.686965

**Authors:** Luz P. Blanco, Binod Regmi, Carmelo Carmona-Rivera, Yudong Liu, Xiantao Wang, Philip M. Carlucci, Monica M. Jackson, Zerai Manna, Sarfaraz Hasni, Markus Hafner, Hong-Wei Sun, Mariana J. Kaplan

## Abstract

**Objective:** Neutrophil dysfunction is a hallmark of systemic lupus erythematosus (SLE), but its molecular basis remains unclear. This study explores transcriptional and post-transcriptional changes in low-density granulocytes (LDGs), a proinflammatory neutrophil subset expanded in SLE, focusing on NADPH oxidase (Nox) function and minor intron splicing.

**Methods:** LDGs and normal-density neutrophils (NDGs) were isolated from SLE patients and healthy controls (HCs). CYBA (P22phox) expression was evaluated at transcript and protein levels. Nox activity was measured using luminol assays. Bulk RNA sequencing and rMATS software were used to assess alternative splicing, particularly of U12-type intron-containing genes.

**Results:** CYBA expression was reduced in SLE LDGs (n=11) compared to SLE NDGs and HCs (n=6), with levels resembling those in chronic granulomatous disease neutrophils. SLE LDGs exhibited impaired Nox activity (n=7 SLE, n=12 HC). CYBA is a U12 intron-containing gene, and transcriptomic analysis revealed broad downregulation of this gene class in SLE LDGs, suggesting minor spliceosome dysfunction. rMATS analysis showed increased U12-type intron retention and widespread splicing defects— including exon skipping and mutually exclusive exon use—in genes such as *GBP5, MAEA* and *STX10*. These abnormalities were validated in an independent long-read RNA-seq dataset from SLE PBMCs. Importantly, splicing disruptions correlated with disease activity and autoantibody profiles.

**Conclusion:** Impaired U12-dependent splicing may contribute to neutrophil dysfunction in SLE, potentially via defective oxidative burst and altered immune regulation. These findings highlight the minor spliceosome as a novel player in lupus pathogenesis.

## Introduction

The pathogenesis of systemic lupus erythematosus (SLE) remains incompletely understood, particularly regarding the role of aberrant RNA splicing in disease progression (1). Widespread splicing abnormalities have been reported in SLE, though it remains unclear whether these disruptions directly generate autoantigens or arise secondarily from chronic autoimmunity, including the production of autoantibodies against nucleic acids and spliceosomal proteins. Notably, autoantibodies targeting spliceosome proteins have been identified in multiple autoimmune conditions, including SLE and murine models of lupus (2–9).

Splicing is a complex regulatory process required for the generation of mature mRNA. While most human genes are processed by the major U2 spliceosome, a small subset (∼700 genes) require the minor U12 spliceosome machinery. These U12-dependent genes are evolutionarily conserved, show tissue-specific expression, and are essential for key biological functions including stress response, immune regulation, and myeloid cell differentiation (10–24). Dysregulated alternative splicing in SLE affects a broad range of immune genes, including *CD44, CD3*ζ*, CTLA4, CREM, Fas*, and *DUSP22*, among others, thereby altering signaling, apoptosis, and tolerance pathways(25–30)

Neutrophils are central to SLE pathogenesis through the production of reactive oxygen species (ROS) and the release of neutrophil extracellular traps (NETs). A distinct subset of low-density granulocytes (LDGs) is expanded in SLE and displays enhanced NET formation, mitochondrial ROS production and distinct pathogenic phenotype (31–33). However, the molecular mechanisms driving their altered function remain incompletely defined.

The *CYBA* gene encoding p22^phox^, is essential for the stability and activity of multiple NADPH (Nox) complexes(34, 35). Reduced *CYBA* expression destabilizes Nox enzymes and impairs oxidative burst, linking defective redox signaling to immune dysregulation(36). In this study, we identify impaired Nox activity in SLE LDGs, accompanied by reduced *CYBA* expression and evidence of U12 intron retention. Transcriptomic analysis further revealed broad splicing abnormalities in SLE LDGs, particularly affecting U12-type introns. These findings suggest that defective minor spliceosome activity may underlie neutrophil dysfunction and contribute to systemic autoimmunity.

## Materials and methods

### Neutrophils and LDG isolation

Neutrophils and LDGs were isolated from peripheral venous blood of SLE subjects and healthy controls (HC), as described (33, 37). SLE patients met revised American College of Rheumatology criteria for this disease (38) and were enrolled in protocol NIH#94-AR-0066. Healthy control volunteers were recruited at the Blood Bank Clinical Center, NIH. All subjects signed informed consent.

### Western blot, immunofluorescence, and flow cytometry

Western blot was performed as previously described (39). Briefly, total protein (50 ug) was resolved in a 4–12% NuPAGE Bis-Tris gradient gel (Invitrogen). Proteins were immobilized onto a nitrocellulose membrane (Invitrogen) and blocked with 10% bovine serum albumin for 30 minutes at room temperature. The membrane was incubated with rabbit anti-human P22phox rabbit polyclonal (1:500; sc-20781, Santa Cruz Biotechnology) or rabbit polyclonal antibody against MPO (1:1,000; A0398, DAKO) overnight at 4°C. After 3 washes with PBS–Tween, the membrane was probed with secondary IRDye800CW goat anti-rabbit antibody (1:10,000; 926-32211, Li-Cor Biosciences) for 1 hour at room temperature. Proteins were detected using a Li-Cor Biosciences Odyssey Infrared Imaging System, in accordance with the manufacturer’s instructions.

For immunofluorescence, neutrophils were seeded into a coverslip and incubated with P22phox rabbit polyclonal (1:500; sc-20781 Santa Cruz Biotechnology) for 1h at room temperature, washed briefly with PBS and incubated with a secondary anti rabbit Alexa 428 antibody (1:250; A21206, Invitrogen) for 1 h at room temperature. After a brief wash in PBS, the cells were fixed in 4% paraformaldehyde overnight at 4°C. After washing in PBS, the coverslips were mounted for fluorescent microscopy analysis using ProLong Gold (P36930, Invitrogen) and visualized using in a Leica DMI4000 B inverted microscope. Images were acquired using the microscope’s associated software.

For the flow cytometry analysis, neutrophils or PBMCs were stained in FACS buffer (PBS and 2% fetal bovine serum) incubated with 100 μL human TruStain FcX on ice (1:50 dilution) for 10 min. After Fc blocking, the cells were incubated with anti-CD15 antibodies (Biolegend), followed by permeabilization with the eBioscience Foxp3/Transcription Factor Staining Buffer Set (Thermo Fisher Scientific). The cells were then stained with anti-P22phox antibody or isotype control antibody (Santa Cruz Biotechnology), followed by staining with fluorescent-conjugated secondary antibodies. Cells were acquired on a BD LSR-Fortessa or BD FACSCanto. Data were analyzed using FlowJo software.

### Nox activity

Nox activity of NDGs and LDGs was measured detecting chemiluminescence produced by luminol once oxidized by Nox in a plate assay. The activity was induced using either N-formyl-Met-Leu-Phe (fMLF) (4 x 10^6 M) or phorbol 12-myristate 13-acetate (PMA) (400 ng/ml). Luminol protected from light was used at 400 uM. A total of 100,000 cells in 100 ul/ well were aliquoted in HBSS containing 10 mM HEPES pH 7.3-7.4. Cells were preincubated with luminol (50 ul) for 10 min at 37°C and then with the activators (50 ul), and luminescence was measured over time up to 30 min and every minute in a Bi-Tek plate reader. The blank control was media alone and cells without activators. The activity is reported as luminescence units (RLU).

### Total RNA sequence

RNAseq libraries were prepared from 1 µg of total RNA using the NEBnext Ultra Directional RNA Library kit (E7760) after ribosomal RNA depletion with the NEB Next rRNA Depletion kit (E7405). RNA-seq data were generated using Illumina’s HiSeq 3000 system. Raw sequencing data were processed with bcl2fastq v2.17.1 to generate FastQ files. Adapter sequences were removed using Trimgalore v0.4.2. Single-end reads of 50 bases were mapped to the human transcriptome and genome hg19 using TopHat v2.1.1. Gene expression values (RPKM: Reads Per Kilobase exon per Million mapped reads) were calculated using Partek Genomics Suite 6.6, which was also used for principal component analysis (PCA) and analysis of variance (ANOVA). Bam files generated by TopHat were used for downstream alternative RNA splicing analysis.

### Alternative Splicing Analysis

Alternative splicing events were analyzed using rMATS v4.3 (40). Multiple unpaired comparisons were performed, with three to ten replicates in the first group (HC) and two to ten replicates in the second group (SLE-LDG, SLE-NDG, SLE-PBMC). rMATS identifies event-based alternative splicing across five patterns: skipped exon (SE), alternative 5′ splice site (A5SS), alternative 3′ splice site (A3SS), mutually exclusive exons (MXE), and retained intron (RI). Significant events were defined by FDR < 0.2(41). These significant alternative splicing events were categorized into two groups based on whether the associated genes were part of the common spliceosome (U2-type) or the rare spliceosome (U12-type). A proportion test was performed to calculate the estimated z-score and p-value, assessing the relative frequency of events in U2- and U12-type genes compared to the total number of genes annotated as U2- or U12-type.

### Pathway analysis

Statistically significant genes with splicing events were analyzed to obtain the relevant pathway plots using the Enrichr web site (https://maayanlab.cloud/Enrichr/) and Appyter viewer (42–44).

### Statistical analysis

The data in figures 2 and 4 was analyzed and plotted using GraphPad Prism software.

## Results

To investigate neutrophil abnormalities in SLE, we first analyzed the expression of CYBA (P22phox), a component of the Nox complex. Western blot analysis revealed reduced CYBA protein levels in lupus LDGs compared to autologous NDGs or heathy control NDGs. Notably, CYBA expression in SLE LDGs was comparable to levels observed in chronic granulomatous disease (CGD) patients (Figure 1A), a condition associated with severely impaired Nox activity (45). Correspondingly, luminol-based assays demonstrated significantly impaired Nox activity in SLE LDGs, with reduced response to both fMLF and PMA stimulation, in contrast to autologous NDGs or healthy control NDGs (Figure 1B). Reduced CYBA (P22phox) expression in SLE LDGs was corroborated by both immunofluorescence staining (Figure 1C) and flow cytometry analysis (Figure 1D).

**Figure 1:**
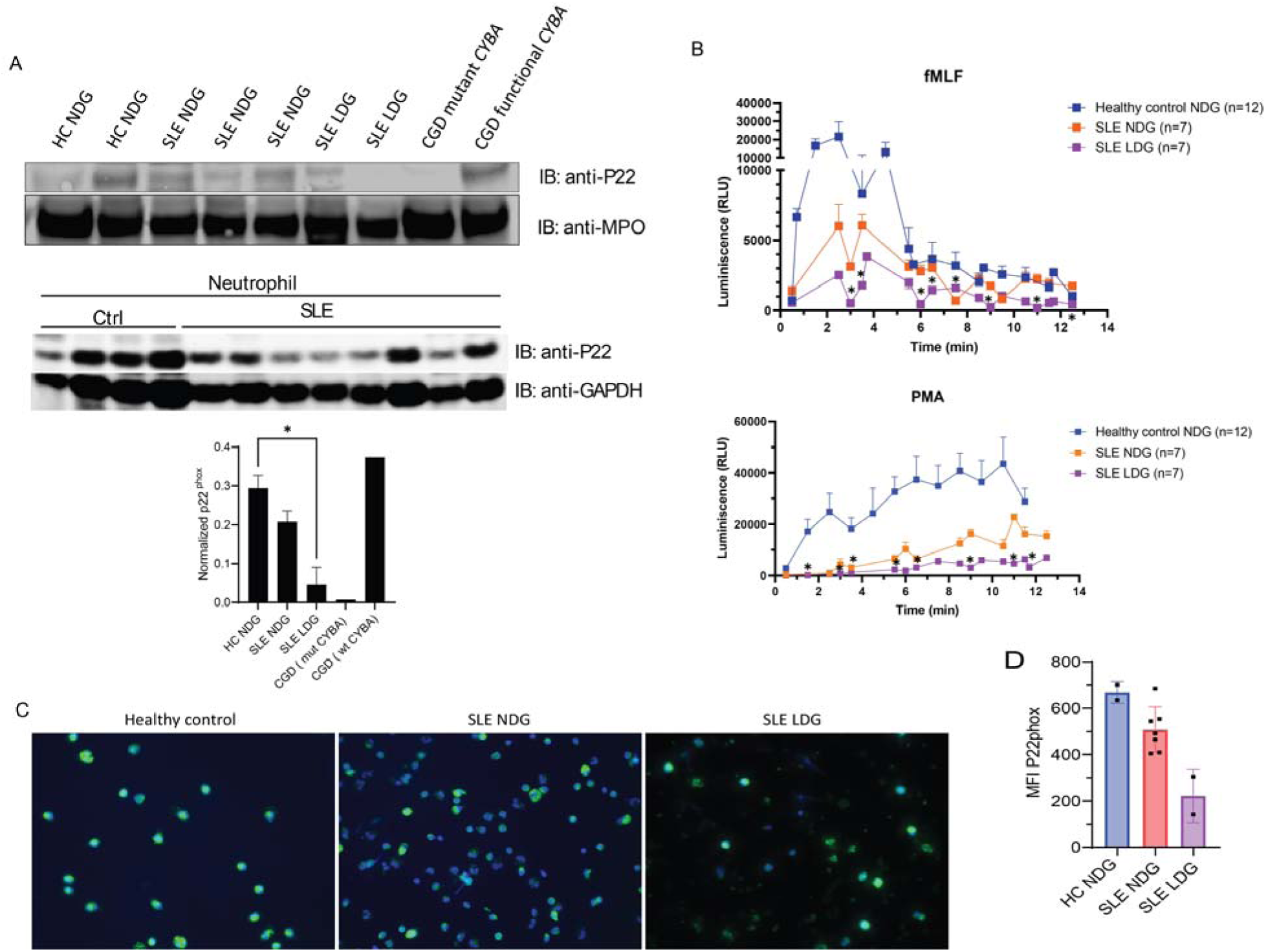
Reduced expression of Nox component CYBA (P22phox) in SLE neutrophils. A) Western blot (upper and middle and respective densitometry analysis (lower) showing the expression of CYBA (P22phox) and myeloperoxidase (MPO, as a house keeping protein) in neutrophils from healthy controls (n=6), SLE (n=11) or CGD (n=2) patients. The top Western gel compares LDGs with autologous NDGs whereas the bottom Western represents NDGs only; * p<0.05. B) Nox activity measured after stimulation with either fMLF (upper graph) or PMA (lower graph) in neutrophils obtained from SLE patients (n=7) or healthy controls (n=12). Asterisks in fMLF indicate statistically significant difference between lupus LDG and NDG, and in PMA significant difference between LDG and healthy control NDG; (*p<0.05), applied Mann-Whitney U-test. C) Images of neutrophils stained with an antibody against P22phox and a secondary antibody (FITC conjugated, green). Nuclei are stained blue with Hoechst (magnification 40x). D) Expression of P22phox in neutrophils measured by flow cytometry. Results represent mean ± SEM. Of SLE patients whose samples were tested for CYBA expression and activity, 83% were anti-ANA and/or anti-ENA positive (50% anti-Ro, 33% anti-Sm, 66% anti-RNP, none anti-La) and mean SLEDAI was 3.6±3.

To explore mechanisms underlying CYBA dysregulation, we analyzed total RNA sequencing data. *CYBA* mRNA expression was lower in SLE LDGs than in NDGs from SLE patients or controls (Figure 2A). Intron retention of *CYBA*, and U12-type minor intron number 2, showed increased retention in SLE LDGs, which inversely correlated with *CYBA* transcript levels (Figure 2B). This was not observed in the housekeeping gene *ACTB*, which showed minimal intron retention and no significant expression difference across cell types (Figure 2C–D). Similarly, the U12-containing gene *PTEN* exhibited reduced expression and increased intron retention in SLE LDGs (Figure 2E–F). Overall, genes harboring U12-type introns were significantly downregulated in SLE LDGs (Figure 2G), consistent with a global impairment of minor spliceosome function.

**Figure 2:**
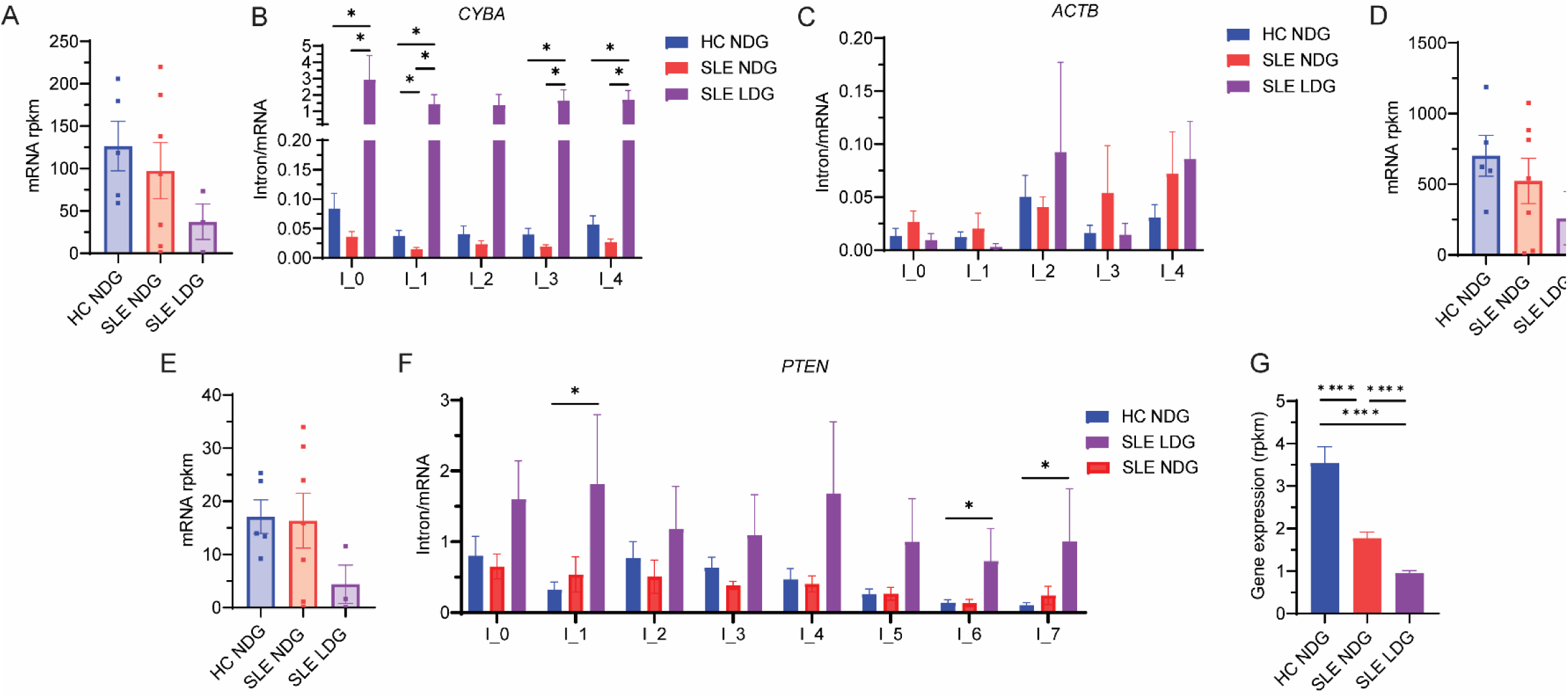
SLE neutrophils display intron retention in genes containing a U12 intron. A) Expression of *CYBA* mRNA in SLE or healthy control neutrophils. B) Intron retention in *CYBA* gene measured in SLE and healthy control neutrophils. C) Intron retention in the *ACTB* housekeeping gene (all U2) and the respective mRNA expression in D). E) *PTEN* mRNA expression and respective intron retention depicted in F). U12 containing genes expression in SLE and healthy control neutrophils in G). Results represent mean ± SEM. *:*p*<0.05, ****:*p*<0.0001. Data analyzed using the Mann-Whitney U-test. Of SLE patients whose samples were sequenced, 86% were ANA positive, 43% were anti-dsDNA positive, 71% were anti-ENA positive (43% anti-Sm positive, 57% anti-Sm/RNP positive, 43% anti-Ro positive and 29% anti-La positive) and mean SLEDAI was 4±3.5.

To test this hypothesis, we used rMATS to analyze alternative splicing events in RNA-seq data from SLE and control neutrophils. Consistent with prior reports, neutrophils from SLE patients (both LDGs and NDGs) exhibited an increased frequency of splicing anomalies. Notably, these disruptions were significantly enriched in genes containing U12-type introns compared to those with U2-type introns (Table 1). Similar splicing patterns were observed in a publicly available long-read RNA-seq dataset of peripheral blood mononuclear cells (PBMCs) from SLE patients (46) suggesting that U12-type splicing dysregulation in SLE may be a systemic feature extending beyond the neutrophil compartment.

**Table 1:** Splicing events are significantly enhanced in U12 genes and in SLE derived cells.

| Cells and comparisons | Exclusive U2 containing genes |  | Exclusive U12 containing genes |  | Number of samples | z-score (H1: P2>P1) | p-value |
| --- | --- | --- | --- | --- | --- | --- | --- |
|  | Total | Number of splicing events | Total | Number of splicing events |  |  |  |
| HC NDG versus SLE LDG | 267 | 307 | 8 | 10 | 6 | 1.9 | 0.030700 |
| SLE NDG versus SLE LDG | 366 | 441 | 14 | 14 | 6 | 3.4 | 0.000290 |
| HC NDG versus SLE NDG | 366 | 427 | 15 | 15 | 5 | 3.8 | 0.000060 |
| HC PBMC versus SLE PBMC | 6,179 | 17,876 | 301 | 827 | 10 | 22.3 | 0.000000 |

The significant splicing events ( $p < 0.05$ ) were filtered (column 3 and column 5) and proportions (P1 and P2) of exclusive number of U2 and U12 containing genes were calculated (column 2 and column 4).
We tested the alternative hypothesis (H1:P2>P1). The Z-score is presented in column 7 and p-value of the scores are presented in column 8. Here, P2 (U12 gene proportion) and P1 (U2 gene proportion)
are the proportion of genes involved in splicing to the total number of genes for each type in the GTF file used for annotation. Abbreviations: HC: healthy control, SLE: systemic lupus erythematosus,
NDG: normal dense neutrophils, LDG: low density neutrophils, PBMC: peripheral blood mononuclear cells.

rMATS analysis identified five categories of aberrant splicing: skipped exons (SE), mutually exclusive exons (MXE), intron retention (RI), and alternative 3’ and 5’ splice sites (A3SS, A5SS). Genes with significant splicing alterations are listed in Table 2 and Supplementary Data (S1–S3). Pathway analysis of these genes (Bioplanet 2019, KEGG 2021) revealed enrichment in immune-related pathways, including interferon and interleukin signaling (Figure 3A), chemokine signaling and MAPK/MEK pathways (Figure 3B), and pathways related to splicing regulation (Figure 3C).

**Figure 3.**
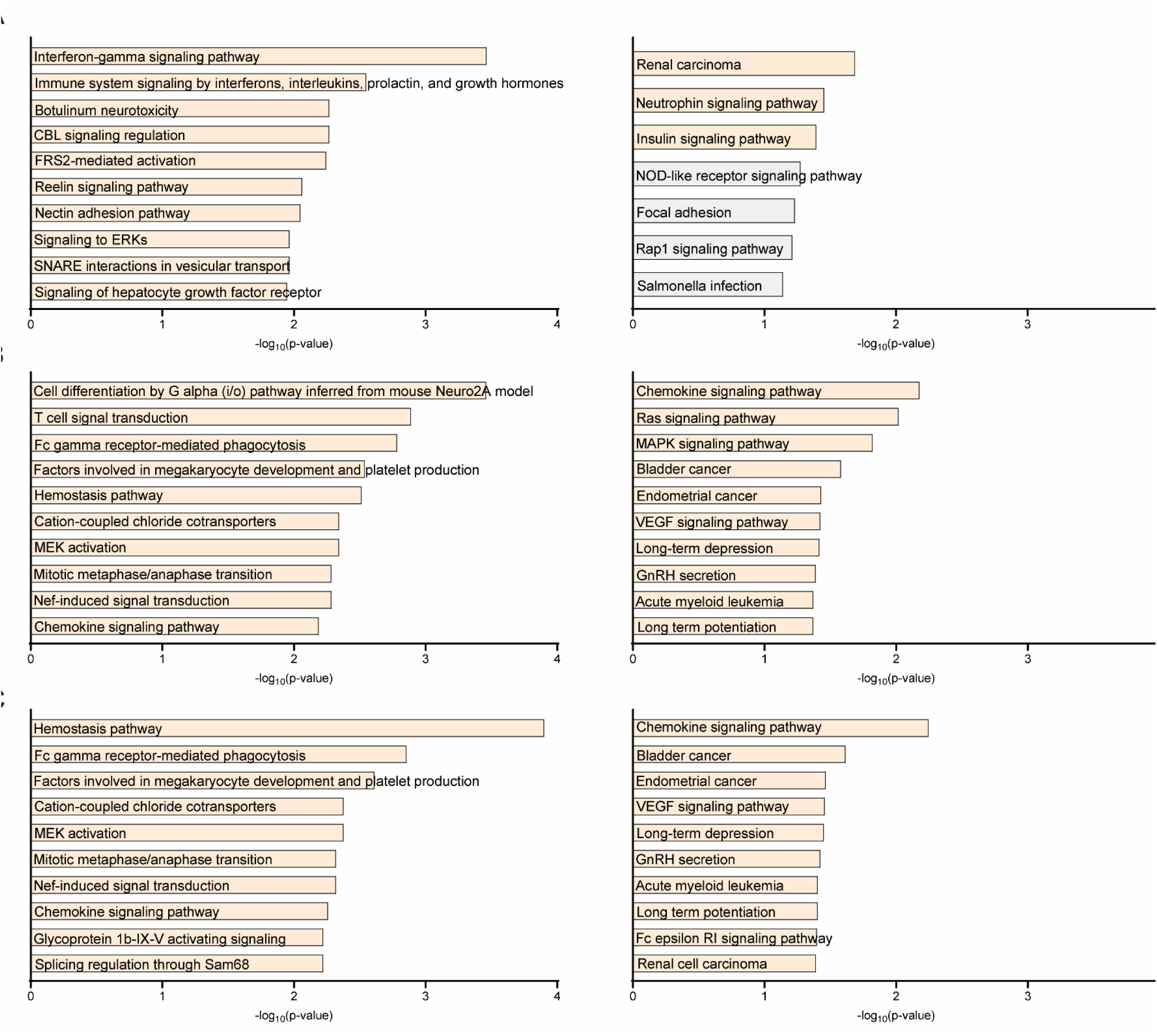
Pathway analysis of significantly regulated genes in SLE neutrophils. Enrichment analysis of differentially expressed or spliced genes was performed using Bioplanet 2019 and KEGG 2021 (Human) pathway databases via the Enrichr platform (https://maayanlab.cloud/Enrichr/) and Appyter viewer. Results are shown for the following comparisons: (A) Healthy control NDGs vs. SLE LDGs; (B) SLE LDGs vs. SLE NDGs, (C) Healthy control NDGs vs. SLE NDGs (**Table 2 for full list).**

**Table 2:** Significant (FDR<0.2) U12 intron containing genes with their respective splicing event.

| HC NDG versus SLE LDG |  |  |  | SLE LDG versus SLE NDG |  |  |  | HC NDG versus SLE NDG |  |  |  |
| --- | --- | --- | --- | --- | --- | --- | --- | --- | --- | --- | --- |
| SE | RI | MXE | A3SS | SE | RI | MXE | A3SS | SE | RI | MXE | A3SS |
| MAEA | (STX10) | GBP5 | None | DISC1 | None | MAEA | (SRPK1) | DISC1 | None | DOCK2 | (SRPK1) |
| (SLC12A9) |  | MAEA |  | DOCK2 |  | RASGRP4 |  | DOCK2 |  | (WDR1) |  |
|  |  | (MAEA) |  | DOCK5 |  |  |  | DOCK5 |  |  |  |
|  |  |  |  | EXOSC2 |  |  |  | EXOSC2 |  |  |  |
|  |  |  |  | GAK |  |  |  | GAK |  |  |  |
|  |  |  |  | GBP5 |  |  |  | GBP5 |  |  |  |
|  |  |  |  | RAF1 |  |  |  | RAF1 |  |  |  |
|  |  |  |  | SBNO2 |  |  |  | SBNO2 |  |  |  |
|  |  |  |  | SLC12A4 |  |  |  | SLC12A4 |  |  |  |
|  |  |  |  | SMC3 |  |  |  | SMC3 |  |  |  |
|  |  |  |  | (SRPK1) |  |  |  | (SRPK1) |  |  |  |

Genes in parenthesis: the splicing event is more predominantly found in the second cell type used for comparison.
Abbreviations: SE: skipped exon, RI: retained intron, MXE: mutually exclusive exons, A3SS: alternative 3' splice site.

Beyond CYBA, we identified splicing disruptions in other components of the Nox complex (*CYBB*, *NCF1, NCF2*, and *NCF4*) across datasets (Supplementary Data S1– S4). Western blot and flow cytometry analyses confirmed reduced, although not statistically significant, levels of CYBB (P47phox) protein in SLE LDGs compared to SLE NDGs and healthy controls (Supplementary Figure S1).

Finally, we investigated the clinical relevance of aberrant splicing events in SLE neutrophils. Splicing anomalies in LDGs showed the strongest associations with clinical disease activity, as measured by SLEDAI score, and with key autoantibody profiles, including anti-dsDNA and anti-Sm (Figure 4A; Supplementary Data S5). Splicing alterations in NDGs were more closely associated with the presence of total anti-ENA and of anti-SmRNP antibodies but did not correlate with disease activity scores (Figure 4B). These findings indicate that splicing dysregulation in LDGs may play a particularly important functional and clinical role in SLE.

**Figure 4.**
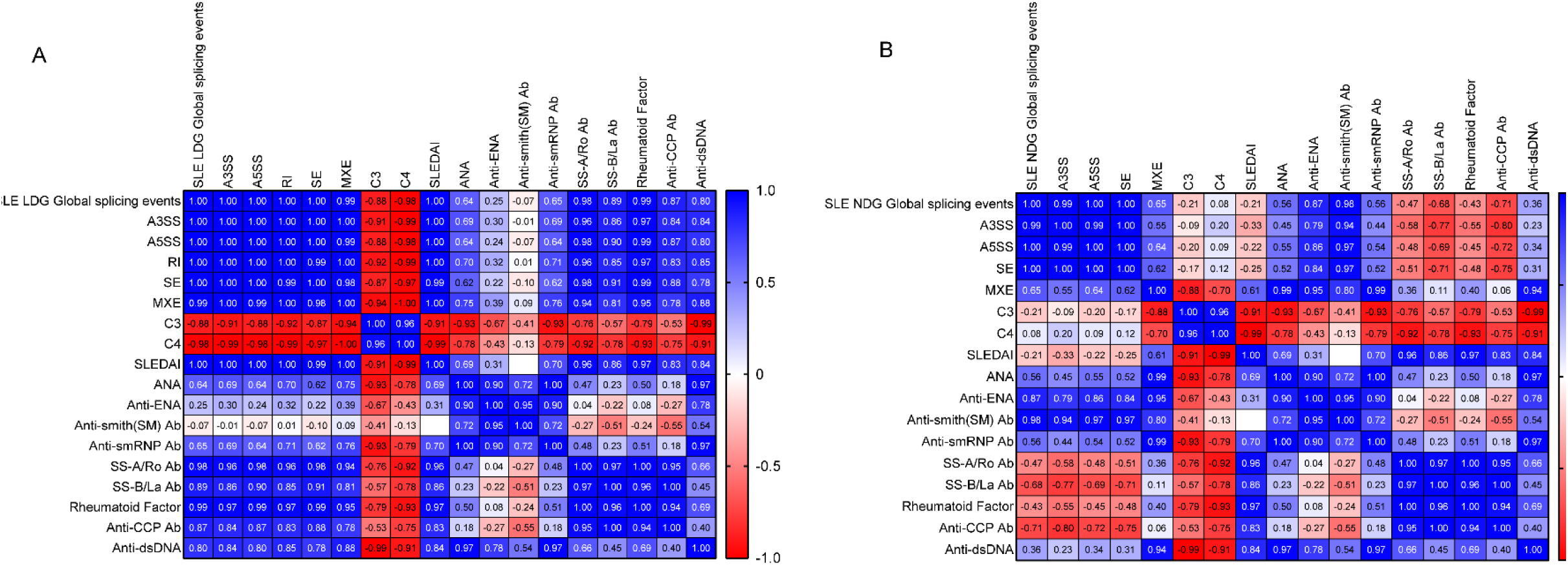
Correlation between clinical and laboratory parameters and splicing events in SLE neutrophils. Pearson correlation coefficients between clinical and laboratory measures and splicing event types in: (A) SLE LDGs vs. healthy control (HC) NDGs and (B) SLE NDGs vs. HC NDGs. Splicing event types include: A3SS: alternative 3′ splice site; A5SS: alternative 5′ splice site; RI: intron retention; SE: exon skipping; MXE: mutually exclusive exon usage. Parameters analyzed: C3: complement component 3; C4: complement component 4; SLEDAI: SLE Disease Activity Index; ANA: antinuclear antibodies; Anti-ENA: anti-extractable nuclear antigens; Anti-SmRNP: anti-Smith ribonucleoprotein antibodies; SS-A/Ro and SS-B/La: autoantibodies targeting Ro and La proteins, respectively; Anti-CCP: anti-cyclic citrullinated peptide antibodies.

Taken together, our results demonstrate widespread alternative splicing abnormalities in SLE neutrophils, with a specific enrichment of defects in U12-type introns. The pronounced splicing disruptions observed in LDGs, especially in genes relevant to immune regulation and oxidative function, may contribute to their pathogenic role and implicate minor spliceosome dysfunction as a mechanistic contributor to SLE pathogenesis.

## Discussion

Our study reveals that neutrophils from SLE patients, especially LDGs, exhibit pervasive splicing abnormalities, with a striking enrichment of aberrant splicing in genes containing U12-type minor introns. These splicing defects likely have profound functional consequences, contributing to altered neutrophil phenotypes and promoting SLE pathogenesis.

Disruptions in U12 intron splicing may impair neutrophil function through multiple mechanisms. First, defective splicing could generate aberrant transcripts and dysfunctional proteins, potentially triggering type I interferon responses or creating neoantigens that drive autoantibody production. This is particularly relevant for LDGs, a subset already characterized by proinflammatory activity and spontaneous, dysregulated NET formation (32).

Among the altered splicing events, mutually exclusive exon usage, a highly conserved and tightly regulated mechanism across tissues (47–49), was particularly enriched in SLE LDGs. For example, *GBP5*, which encodes an interferon-inducible GTPase involved in inflammatory responses and lupus nephritis progression in mice (50, 51) exhibited both mutually exclusive exon usage and exon skipping in lupus neutrophils compared to healthy controls. GBP5+ neutrophils with elevated interferon signatures have also been identified in inflamed human tissues, such as nasal polyps, indicating broader relevance beyond lupus (52).

In NDGs, splicing disruptions were detected in *SRPK1*, a splicing kinase involved in neutrophil development and linked to myelodysplastic syndromes (53). Additionally, several *MAPK* family members, many of which contain U12 introns and play essential roles in stress signaling, were affected (Supplementary Data S1-4). Importantly, U12 intron-containing genes are generally less efficiently spliced due to the instability of U6atac, a key snRNA in the minor spliceosome (54, 55). Notably, *MAPK14* (p38), itself U12-regulated, is critical for stabilizing U6atac under stress. This suggests a feedback loop where impaired p38 expression may further destabilize U12 splicing, compounding splicing dysfunction in SLE neutrophils. This mechanism may also operate in other immune cells, as indicated by the U12 splicing abnormalities detected in SLE PBMCs. (Supplementary Data S4).

Additional support for a splicing-driven dysfunction in neutrophils comes from altered splicing in *DOCK2*, a Rac activator essential for actin cytoskeleton dynamics and oxidative burst (56). Combined with multiple splicing events detected in Nox components beyond *CYBA*, these findings offer a putative mechanistic explanation for the reduced respiratory burst observed in SLE LDGs.

Our data reinforce the distinctiveness of the LDG subset and implicate it in SLE pathogenesis through both compromised Nox function and pervasive U12 splicing defects. This aligns with genetic studies linking polymorphisms in Nox genes to elevated SLE risk (57–59). Importantly, U12 splicing anomalies in LDGs associate with disease activity and specific autoantibody profiles, underscoring a possible role for splicing errors in generating autoantigens and amplifying autoimmune responses in SLE.

While we observed a trend toward reduced CYBB protein levels in SLE LDGs compared to NDGs and healthy controls, this difference did not reach statistical significance. Nevertheless, this pattern complements the broader Nox dysregulation observed. CYBB and CYBA (p22phox) are critical Nox subunits responsible for ROS generation during the oxidative burst. The concurrent reduction of these components in LDGs suggests a coordinated impairment of complex assembly or stability, potentially contributing to the diminished oxidative burst capacity. Since *CYBB* lacks a U12-type intron, its reduced expression likely reflects additional mechanisms beyond minor spliceosome dysfunction, such as altered transcriptional regulation, enhanced protein degradation, retrotransposable genetic elements causing mutations, or LDG-specific epigenetic modifications (18, 60, 61). Collectively, these findings highlight multifactorial defects in the oxidative machinery of LDGs that may underlie their heightened inflammatory and pathogenic potential.

A plausible driver of the widespread splicing abnormalities, especially those involving U12 introns, is chronic cellular stress stemming from persistent systemic inflammation and oxidative imbalance. LDGs are known to produce elevated mitochondrial ROS (33), which could impair spliceosome components or their regulatory kinases. The intrinsic instability of *U6atac* snRNA further sensitizes U12 splicing to stress-induced disruption. Genetic factors, such as SNPs in spliceosome-related genes or regulatory kinases like SRPK1 and MAPK14/p38, may exacerbate splicing defects (62, 63). Moreover, autoantibodies targeting spliceosomal proteins, commonly present in SLE patients, might directly impair splicing machinery, increasing transcriptomic noise and producing aberrant RNA species. These combined disruptions could promote novel autoantigen formation, perpetuating autoimmunity and disease progression.

Although both LDGs and NDGs from SLE patients exhibited increased retention of U12-type introns, the decrease in *CYBA* expression was specific to LDGs. This selective downregulation suggests that U12 splicing defects alone do not fully explain altered gene expression in SLE neutrophils. Instead, cell-intrinsic factors unique to LDGs, such as altered maturation, epigenetic remodeling, and activation states (32), likely influence transcript stability, translation efficiency, or protein turnover. For example, *CYBA* transcripts may be more susceptible to degradation of mis-spliced RNA or differential regulation by RNA-binding proteins within LDGs. These findings support the concept that the functional impact of U12 splicing dysregulation is context-dependent, preferentially disrupting gene networks critical for oxidative burst and immune regulation in distinct neutrophil subsets.

While our study provides novel insights into U12 intron splicing defects and neutrophil dysfunction in SLE, several limitations warrant consideration. First, the sample size was modest, which limits statistical power and the robustness of some clinical correlations. This includes uneven group sizes in RNA-seq comparisons, which increase the risk of false discovery. Larger, well-stratified cohorts will be required to confirm these preliminary findings and extend them to broader patient populations, including individuals with CGD. Second, while transcriptomic analysis revealed widespread splicing changes, functional validation of specific splice variants, such as their effects on protein expression and enzymatic activity, was only partially addressed. Future studies should directly define how these splicing alterations affect neutrophil biology and immune responses. Finally, although we observed correlations with clinical features, causal relationships remain unproven, and our focus on neutrophils does not exclude the possibility that U12 splicing defects may extend to other immune cell types in SLE, as suggested by PBMC analyses.

In summary, our findings highlight disrupted U12 intron splicing as a previously underappreciated contributor to neutrophil dysfunction in SLE, particularly within the pathogenic LDG subset. This study broadens our understanding of lupus pathogenesis and opens new avenues for developing therapeutic strategies that target RNA splicing pathways.

## Acknowledgment

This work utilized the computational resources of the NIH HPC Biowulf cluster (https://hpc.nih.gov). This study was supported by the Intramural Research Program at NIAMS (ZIA AR041199).

This research was supported [in part] by the Intramural Research Program of the National Institutes of Health (NIH). The contributions of the NIH author(s) were made as part of their official duties as NIH federal employees, are in compliance with agency policy requirements, and are considered Works of the United States Government. However, the findings and conclusions presented in this paper are those of the author(s) and do not necessarily reflect the views of the NIH or the U.S. Department of Health and Human Services.

## Author Contributions

All authors were involved in revising the manuscript for important intellectual content, and all authors approved the final version to be published. Dr Blanco wrote the original version of this manuscript. Drs. Kaplan and Sun have full access to all the data in the study and the manuscript responsibility for the integrity of the data and the accuracy of the data analysis.

Study conception and design

Blanco, Sun, Kaplan.

Acquisition of data

Blanco, Remi, Liu, Wang, Carlucci, Jackson, Carmona-Rivera, Hasni, Hafner, Sun.

Analysis and of data

Blanco, Remi, Sun.

